# The construction and deconstruction of sub-optimal preferences through range-adapting reinforcement learning

**DOI:** 10.1101/2020.07.28.224642

**Authors:** Sophie Bavard, Aldo Rustichini, Stefano Palminteri

**Affiliations:** Laboratoire de Neurosciences Cognitives et Computationnelles, Institut National de la Santé et Recherche Médicale, 29 rue d’Ulm 75005 Paris, FR; Ecole normale supérieure, 29 rue d’Ulm 75005 Paris, FR; Université de Recherche Paris Sciences et Lettres, 60 rue Mazarine 75006 Paris, FR; University of Minnesota, 1925 4th Street South 4-101, Hanson Hall, Minneapolis, USA

## Abstract

Converging evidence suggests that economic values are rescaled as a function of the range of the available options. Critically, although locally adaptive, range adaptation has been shown to lead to suboptimal choices. This is particularly striking in reinforcement learning (RL) situations when options are extrapolated from their original context. Range adaptation can be seen as the result of an adaptive coding process aiming at increasing the signal-to-noise ratio. However, this hypothesis leads to a counter-intuitive prediction: decreasing outcome uncertainty should increase range adaptation and, consequently, extrapolation errors. Here, we tested the paradoxical relation between range adaptation and performance in a large sample of subjects performing variants of a RL task, where we manipulated task difficulty. Results confirmed that range adaptation induces systematic extrapolation errors and is stronger when decreasing outcome uncertainty. Finally, we propose a range-adapting model and show that it is able to parsimoniously capture all the observed results.

## Introduction

In the famous Ebbinghaus illusion, two circles of identical size are placed near to each other, and larger circles surround one, while smaller circles surround the other. As a result, the central circle surrounded by larger circles appears smaller than the central circle surrounded by smaller circles, as if its subjective estimation of size was affected by the surroundings. Beyond perceptual decision-making, wealth of evidence in neuroscience and in economics suggests that the subjective economic value of one option is not estimated in isolation, but is highly dependent of the context in which the options are presented [1,2]. The vast majority of neuroeconomic studies of context-dependent valuation in humans considered situations where subjective values are triggered by explicit cues, that is stimuli whose value can be directly inferred, such as lotteries or snacks [3–5]. However, in a series of recent papers, we and other groups demonstrated that contextual adjustments also permeate reinforcement learning situations, i.e., when option values have to be inferred from the history of past outcomes [6–8]. We showed that an option, whose small objective value (7.5c) is learned in a context of smaller outcomes, is preferred to an option whose objective value (25c) is learned in a context of bigger outcomes, thus providing an economic equivalent of the Ebbinghaus illusion. Similar observations in birds suggest that this is a feature of decision-making broadly shared across vertebrates [9,10].

Although (as illustrated in the previous example) value context-dependence may lead to suboptimal decisions, it could be normatively understood as an adaptive process aimed at rescaling the behavioral response as a function of the range of the available options. Specifically, it could be seen as the result of an adaptive coding process aiming at increasing the signal-to-noise ratio by a system (the brain) constrained by the fact that behavioral variables have to be encoded by finite firing rates. In other terms, range adaptation would be a consequence of how the system adjusts and optimizes the function associating the firing rate to the objective value to put its slope its maximum for each context [11,12]. The hypothesis that value context-dependence in reinforcement learning is the consequence of adaptive coding entails a counter-intuitive prediction. Specifically, if range adaptation is a direct consequence of the way the brain encodes external information (i.e., is an automatic process), factors that reduce the uncertainty about context-level variables (e.g., the maximum and minimum outcome) should decrease the deviation between objective values (context-independent or absolute) and subjective values (context-dependent or relative). This prediction is in striking contrast with the virtually universally shared intuition that making a decision-problem easier (by facilitating the identification of its variables should, if anything, lead to more accurate and objective internal representations. In the present study, we aim at testing this hypothesis, while concomitantly gaining a better understanding of range adaptation at the computational level. To empirically test this hypothesis, we opted for a reinforcement learning paradigm that, compared to different set-ups (such as lotteries), has the advantage of being straightforwardly translatable to animal and artificial intelligence research [13]. Building on previous research, our task featured a learning phase and a transfer phase [6]. In the learning phase, participants had to determine by trial-and-error the most favorable option in four fixed pairs of options (contexts), with different outcome ranges. In transfer phase, the original options were rearranged, thus creating new contexts. This set up allowed us to quantify learning (or acquisition) errors during the first phase, and extrapolation (or transfer) errors during the second phase. Crucially, the task contexts were designed such that the correct responses in the transfer phase presented an overall higher expected value. We declined this paradigm in eight different versions where we manipulated the task difficulty in complementary ways. First, half of the experiments featured complete feedback information, meaning that participants were informed about the outcome of the forgone option. This manipulation intrinsically reduces uncertainty about the option values and has been repeatedly shown to improve learning performance [8, 14]. Second, half of the experiments featured a blocked (instead of interleaved design), meaning that all the trials featuring one context were presented in a row. This manipulation extrinsically reduces uncertainty about the option values by reducing working memory demand and has also been shown to improve learning performance [15]. Finally, in half of the experiments feedback was also provided in the transfer phase, thus allowing to assess if and how the values learned during the learning phase can be revised. Behavioral analyses backed up our prediction and indicate that learning errors and extrapolation errors are largely dissociable. Critically (and paradoxically), transfer phase performance was lower when the learning phase was easier. Accordingly, the estimated deviation between the objective values and the subjective values was augmented in the complete feedback and block design tasks. Only adding complete feedback in the transfer phase allowed correcting the deviation.

To complement choice rate analysis, we developed a computational model that implements range adaption by tracking the maximum possible reward in each learning context. Model simulations and out-of-sample validation parsimoniously captured performance in the learning and the transfer phase, including the range adaptation-induced suboptimal preferences. Model simulations also allowed us to rule out alternative interpretations of our results that could come from two prominent psychological and economic theories: habit formation and risk aversion [16,17].

## Results

### Experimental protocol

We designed a series of learning and decision-making experiments involving variants of a main task. The main task was composed of two phases: the learning and the transfer phase. During the learning phase, participants were presented with eight abstract pictures, organized in four stable choice contexts. In the learning phase, each choice context featured only two possible outcomes: either 10pt/0pt or 1pt/0pt. The outcomes were probabilistic (75% or 25%) and we labeled the choices contexts as a function of the difference in expected value between the most and the least rewarding option: ΔEV=5 and ΔEV=0.5 (**Figure 1A**). In the subsequent transfer phase, the eight options were re-arranged into new choice contexts, where options associated with 10pt were compared to options associated with 1pt (see [7, 10] for similar designs in humans and starlings). The resulting new four contexts were labeled ΔEV=7.25, ΔEV=6.75, ΔEV=2.25, ΔEV=1.75 (**Figure 1B**).

**Figure 1:**
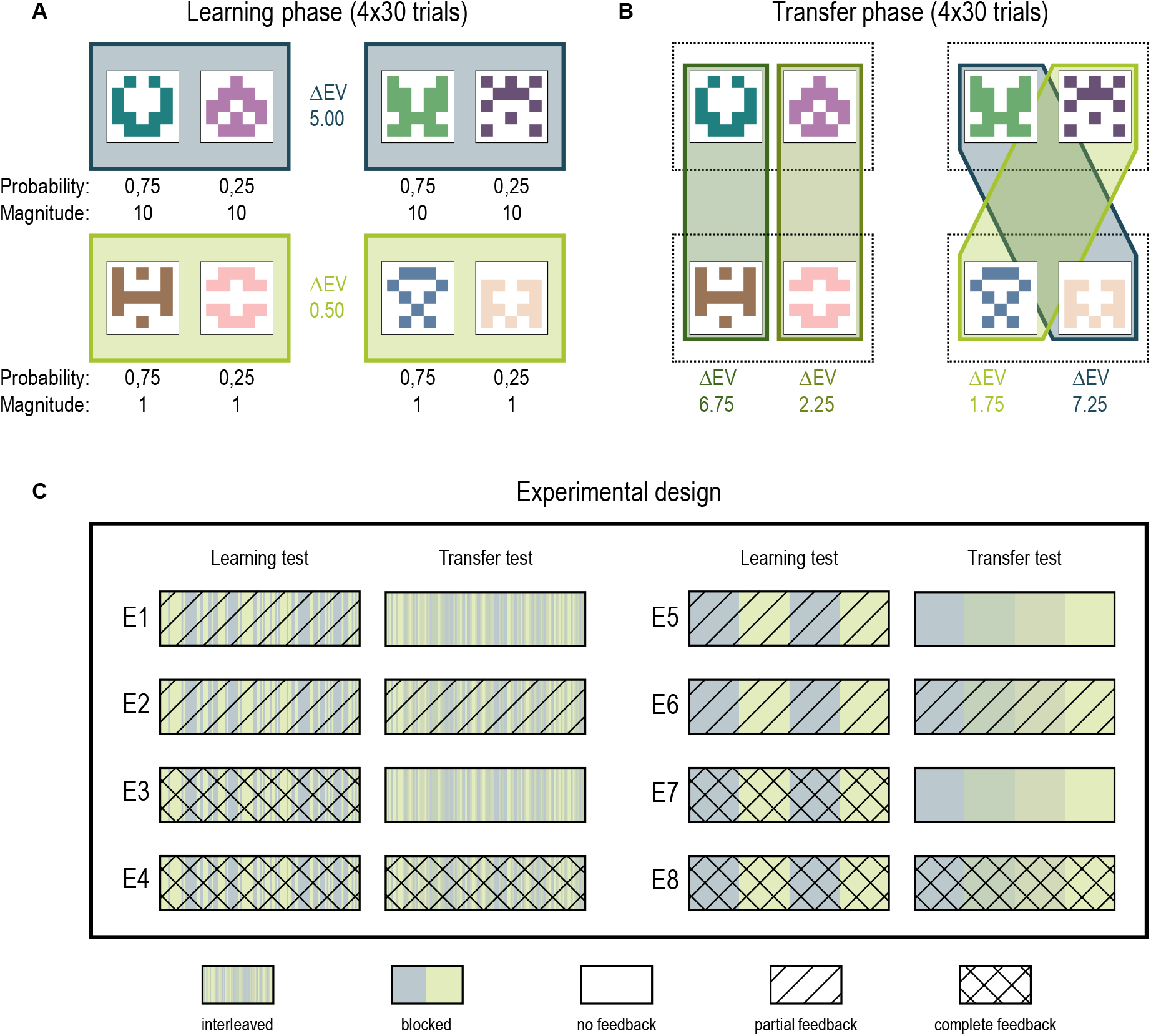
Experimental design. **(A)** Choice contexts in the learning phase. During the learning phase subjects were presented with four choice contexts, including high magnitude (ΔEV = 5.0 contexts) and low magnitude (ΔEV = 0.5 contexts). **(B)** Choice contexts in the transfer phase. The four options were re-arranged into four new choice contexts, each involving both the 1pt and the 10pt outcome. **(C)** Experimental design. The eight experiments varied in the temporal arrangement of choice contexts (interleaved or blocked) and the quantity of feedback in the learning phase (partial or complete) and the transfer phase (without or with).

In our between-subjects study, we developed eight different variants of the main paradigm where we manipulated whether we provided trial-by-trial feedback in the transfer phase (with / without), the quantity of information provided at feedback (partial / complete) and the temporal structure of choice contexts presentation (interleaved / blocked) (**Figure 1C**). All the experiments reported in the main text were conducted online (N=100 subjects in each version of the experiment); we report in the supplementary information the results concerning a similar experiment realized in the lab (see **Supplementary Materials**).

### Overall correct response rate

The main dependent variable in our study was the correct response rate, i.e., the proportion of expected value-maximizing choices in the learning and the transfer phase (crucially our task design allowed to identify an expected value-maximizing choice in all choice contexts). In the learning phase, the average correct response rate was higher than chance level (0.5), indicating overall significant and robust reinforcement learning effect (0.69 ± 0.16, *t*(799) = 32.49, *p* < .0001; **Figure 2A-B**). Replicating previous findings, in the learning phase, we also observed a moderate but significant effect of the choice contexts, where the correct choice rate was higher in the ΔEV=5.0 compared to the ΔEV=0.5 contexts (0.71 ± 0.18 vs 0.67 ± 0.18; *t*(799) = 6.81, *p* < .0001, *d* = 0.24; **Figure 2C**) [6]. Correct response rate was also higher than chance in the transfer phase (0.62 ± 0.17, *t*(799) = 20.29, *p* < .0001, *d* = 0.72, **Figure 2D-E**), where it was also strongly modulated by the choice context (*F*(2.84, 2250.66) = 271.68, *p* < .0001, 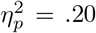, Huynh–Feldt corrected). In the transfer phase, the latter ΔEV=1.75 choice context is of particular interest, since the expected value maximizing option was the least favorable option of a ΔEV=5.0 context in the learning phase, and, conversely, the expected value minimizing option was the most favorable option of a ΔEV=0.5 context of the learning phase. Therefore, absolute versus relative values learning predict opposite preferences in this choice context. Crucially, in the ΔEV=1.75 context, we found that participants’ average correct choice rate was significantly below chance level (0.42 ± 0.30, *t*(799) = −7.25, *p* < .0001, *d* = −0.26; **Figure 2F**), thus demonstrating that participants express suboptimal (i.e., expected value minimizing) preferences in this context.

**Figure 2:**
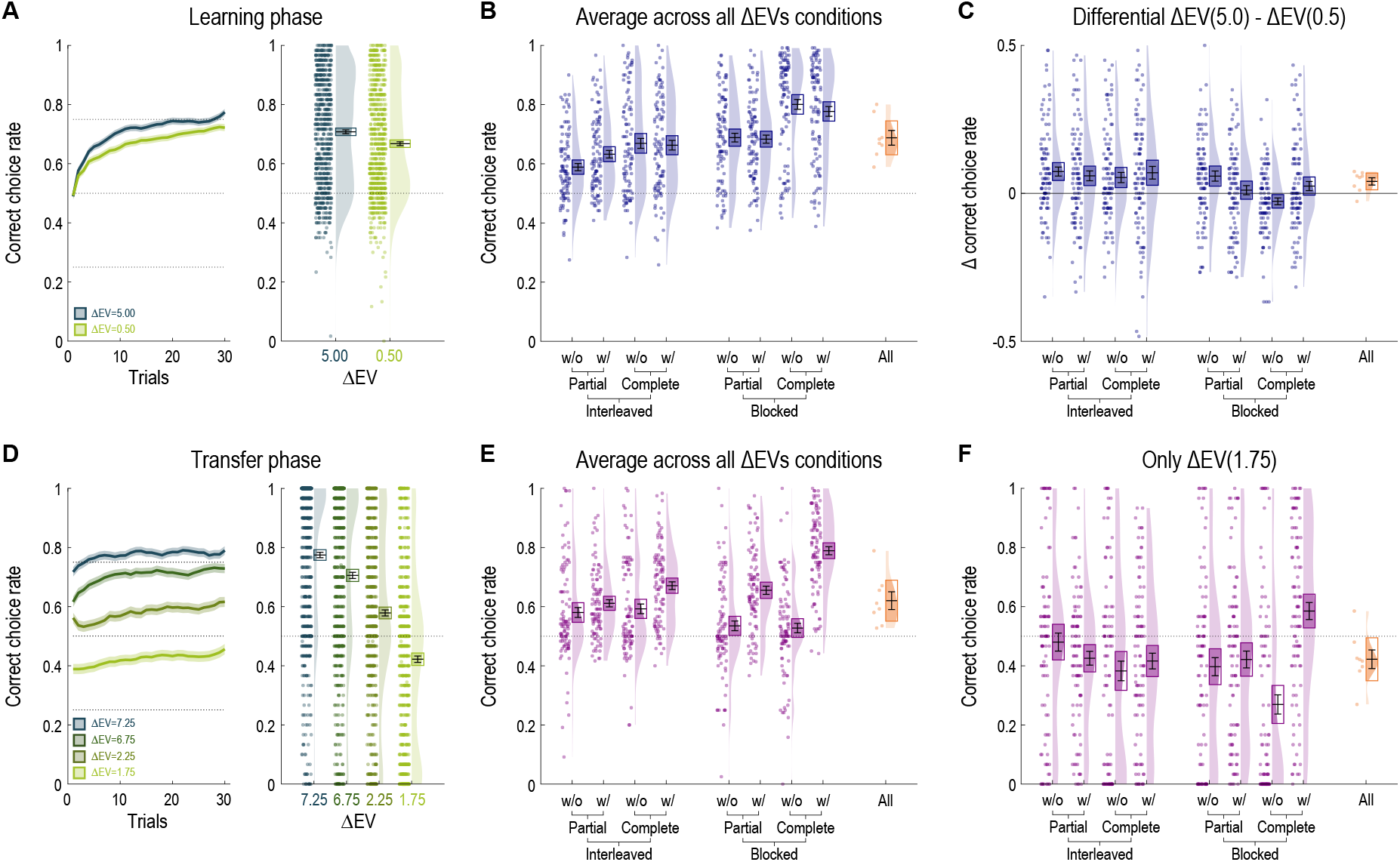
Behavioral results. **(A)** Correct choice rate in the learning phase as a function of the choice context (ΔEV=5.0 or ΔEV=0.5). Leftmost panel: learning curves; rightmost panel: average across all trials. **(B)** Average correct response rate in the learning phase per experiment (in blue: N=800 participants) and meta-analytical (in orange: N=8 experiments). **(C)** Difference in correct choice rate between the ΔEV=5.0 and the ΔEV=0.5 contexts per experiment (in blue: N=800 participants) and meta-analytical (in orange: N=8 experiments). **(D)** Correct choice rate in the transfer phase as a function of the choice context (ΔEV=7.25, ΔEV=6.75, ΔEV=2.25 or ΔEV=1.75). Leftmost panel: learning curves; rightmost panel: average across all trials. **(E)** Average correct response rate in the transfer phase per experiment (in pink: N=800 participants) and meta-analytical (in orange: N=8 experiments). **(F)** Correct choice rate for the ΔEV=1.75 context only (in pink: N=800 participants) and meta-analytical (in orange: N=8 experiments). In all panels: points indicate individual average, areas indicate probability density function, boxes indicate 95% confidence interval and errors bars indicate s.e.m.

### Between-experiments comparisons: learning phase

In this section we analyze the correct response rate as a function of the experimental factors manipulated across the eight experiments (the quantity of provided information, that could be either partial or complete; the temporal structure of choice contexts presentation, that could be blocked or interleaved; and whether feedback was provided in the transfer phase). In the main text we report the significant results, but please see **Tables 1 and 2** for all results and effect sizes.

**Table 1:** Statistical effects of the ANOVA on the choice rate as a function of task factors. LF: learning feedback (partial/complete). TF: transfer feedback (without/with). BE: block effect (interleaved/blocked). PE: phase effect (learning/transfer). DFn: degrees of freedom numerator. DFn: degrees of freedom denominator. \**p* <.05, \*\**p* <.01, \*\*\**p* <.001

|  |  |  | Learning performance |  |  |  | Transfer performance |  |  |  | Overall performance |  |  |  |
| --- | --- | --- | --- | --- | --- | --- | --- | --- | --- | --- | --- | --- | --- | --- |
| | DFn | DFd | F-val | | P-val | $\eta_p^2$ | F-val | | P-val | $\eta_p^2$ | F-val | | P-val | $\eta_p^2$ |
| **LF - Learning feedback** | 1 | 792 | 55,57 | *** | 2,36E-13 | 0,18 | 22,36 | *** | 2,68E-06 | 0,01 | 61,68 | *** | 1,31E-14 | 0,11 |
| **TF - Transfer feedback** | 1 | 792 | 0,04 |  | 0,84 | 0,00 | 137,18 | *** | 2,47E-29 | 0,07 | 58,11 | *** | 7,10E-14 | 0,10 |
| **BE - Block effect** | 1 | 792 | 87,22 | *** | 9,60E-20 | 0,25 | 1,53 |  | 0,22 | 0,00 | 46,82 | *** | 1,56E-11 | 0,08 |
| **PE - Phase effect** | 1 | 792 | - |  | - | - | - |  | - | - | 103,07 | *** | 7,56E-23 | 0,12 |
| **LF x TF** | 1 | 792 | 2,61 |  | 0,1066 | 0,01 | 20,18 | *** | 8,09E-06 | 0,01 | 3,33 |  | 0,0682 | 0,01 |
| **LF x BE** | 1 | 792 | 5,05 | * | 0,0249 | 0,02 | 1,66 |  | 0,1977 | 0,00 | 5,20 | * | 0,0229 | 0,01 |
| **TF x BE** | 1 | 792 | 2,43 |  | 0,1194 | 0,01 | 42,22 | *** | 1,44E-10 | 0,02 | 9,89 | ** | 0,0017 | 0,02 |
| **LF x PE** | 1 | 792 | - |  | - | - | - |  | - | - | 4,97 | * | 0,0261 | 0,01 |
| **TF x PE** | 1 | 792 | - |  | - | - | - |  | - | - | 82,30 | *** | 9,07E-19 | 0,09 |
| **BE x PE** | 1 | 792 | - |  | - | - | - |  | - | - | 42,09 | *** | 1,54E-10 | 0,05 |
| **LF x TF x BE** | 1 | 792 | 0,55 |  | 0,4603 | 0,00 | 5,02 | * | 0,0253 | 0,00 | 3,65 |  | 0,0563 | 0,01 |
| **LF x TF x PE** | 1 | 792 | - |  | - | - | - |  | - | - | 23,37 | *** | 1,61E-06 | 0,03 |
| **LF x BE x PE** | 1 | 792 | - |  | - | - | - |  | - | - | 0,61 |  | 0,4365 | 0,00 |
| **TF x BE x PE** | 1 | 792 | - |  | - | - | - |  | - | - | 40,58 | *** | 3,19E-10 | 0,05 |
| **LF x TF x BE x TE** | 1 | 792 | - |  | - | - | - |  | - | - | 1,39 |  | 0,2396 | 0,00 |

**Table 2:** Participants’ age and correct choice rate as a function of experiments and task factors.

|  | Experiment 1<br>N=100 |  | Experiment 2<br>N=100 |  | Experiment 3<br>N=100 |  | Experiment 4<br>N=100 |  | Experiment 5<br>N=100 |  | Experiment 6<br>N=100 |  | Experiment 7<br>N=100 |  | Experiment 8<br>N=100 |  | Total<br>N=800 |  |
| --- | --- | --- | --- | --- | --- | --- | --- | --- | --- | --- | --- | --- | --- | --- | --- | --- | --- | --- |
|  | mean | std | mean | std | mean | std | mean | std | mean | std | mean | std | mean | std | mean | std | mean | std |
| Age | 30,48 | 10,70 | 27,23 | 8,30 | 32,01 | 10,51 | 31,57 | 9,80 | 33,04 | 10,48 | 28,46 | 10,20 | 28,73 | 9,89 | 28,84 | 9,60 | 30,06 | 10,10 |
| % correct |  |  |  |  |  |  |  |  |  |  |  |  |  |  |  |  |  |  |
| Learning phase | 0,59 | 0,12 | 0,63 | 0,13 | 0,67 | 0,17 | 0,66 | 0,16 | 0,69 | 0,15 | 0,68 | 0,14 | 0,80 | 0,17 | 0,78 | 0,16 | 0,69 | 0,16 |
| ΔEV=5,0 | 0,63 | 0,16 | 0,66 | 0,17 | 0,70 | 0,19 | 0,70 | 0,20 | 0,72 | 0,17 | 0,69 | 0,17 | 0,79 | 0,19 | 0,79 | 0,18 | 0,67 | 0,18 |
| ΔEV=0,5 | 0,55 | 0,13 | 0,60 | 0,14 | 0,64 | 0,19 | 0,63 | 0,19 | 0,66 | 0,17 | 0,68 | 0,14 | 0,81 | 0,17 | 0,76 | 0,18 | 0,71 | 0,18 |
| Transfer phase | 0,58 | 0,17 | 0,61 | 0,12 | 0,59 | 0,16 | 0,67 | 0,13 | 0,54 | 0,16 | 0,66 | 0,14 | 0,53 | 0,16 | 0,79 | 0,14 | 0,62 | 0,17 |
| ΔEV=7,25 | 0,67 | 0,28 | 0,76 | 0,22 | 0,75 | 0,29 | 0,85 | 0,19 | 0,66 | 0,30 | 0,84 | 0,18 | 0,76 | 0,31 | 0,93 | 0,14 | 0,77 | 0,26 |
| ΔEV=6,75 | 0,64 | 0,29 | 0,68 | 0,26 | 0,70 | 0,31 | 0,81 | 0,19 | 0,62 | 0,32 | 0,76 | 0,27 | 0,55 | 0,37 | 0,89 | 0,16 | 0,71 | 0,30 |
| ΔEV=2,25 | 0,54 | 0,27 | 0,58 | 0,19 | 0,54 | 0,34 | 0,61 | 0,28 | 0,47 | 0,32 | 0,60 | 0,18 | 0,54 | 0,36 | 0,76 | 0,22 | 0,58 | 0,29 |
| ΔEV=1,75 | 0,48 | 0,30 | 0,43 | 0,23 | 0,38 | 0,33 | 0,42 | 0,27 | 0,40 | 0,31 | 0,42 | 0,28 | 0,27 | 0,32 | 0,59 | 0,29 | 0,42 | 0,30 |

First, we analyzed the correct choice rate in the learning phase (**Figure 2B**). As expected, increasing feedback information had a significant effect on correct choice rate in the learning phase (*F*(1,792) = 55.57, *p* < .0001, 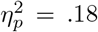); similarly, performance in the block design experiments was significantly higher (*F*(1, 792) = 87.22, *p* < .0001, 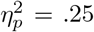). We found a significant interaction between feedback information and task structure, reflecting that the difference of performance between partial and complete feedback was higher in block design (*F*(1, 792) = 5.05, *p* = .02, 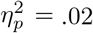). We found no other significant main effect, double or triple interaction (**Table 1**). We also analyzed the difference in performance between the ΔEV=5.0 and ΔEV=0.5 choice contexts across experiments (**Figure 2C**). We found a small but significant effect of temporal structure, the differential being smaller in the blocked compared to interleaved experiments (*F*(1,792) = 7.71, *p* = .006, 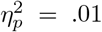), and found no other significant main effect, nor interaction. To sum up, as expected [8, 14, 15], increasing feedback information and clustering the choice contexts had a beneficial effect on correct response rate. Designing the choice contexts in blocks also blunted the difference in performance between the small (ΔEV=0.5) and big (ΔEV=5.0) magnitude contexts.

### Between-experiments comparisons: transfer phase

We then analyzed the correct choice rate in the transfer phase (**Figure 2E**). Unsurprisingly, showing trial-by-trial feedback in the transfer phase led to significantly higher performance (*F*(1, 792) = 137.18, *p* < .0001, 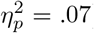). Increasing feedback information from partial to complete also had a significant effect on transfer phase correct choice rate (*F*(1,792) = 22.36, *p* < .0001, 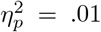).

Interestingly, we found no significant main effect of task structure in the transfer phase (see **Table 1**). We found a significant interaction between feedback and the presence of feedback in the transfer phase, showing that the increase in performance due to the addition of feedback information is higher when both outcomes were displayed during the learning phase (*F*(1,792) = 20.18, *p* < .0001, 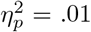). We also found a significant interaction between transfer feedback and task structure, reflecting that the increase in performance due to the addition of feedback information was even higher in block design (*F*(1, 792) = 42.22, *p* < .0001, 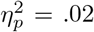). Finally, we found a significant triple interaction between feedback information, the presence of feedback in the transfer phase, and task structure (*F*(1, 792) = 5.02, *p* = . 03, 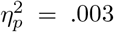). We found no other significant double interaction. We also separately analyzed the correct choice rate in the ΔEV=1.75 context (**Figure 2F**). Overall, the statistical effects presented a similar pattern as the correct choice rate across all conditions (see **Table 2**), indicating that overall correct choice rate and the correct choice rate in the key comparison ΔEV=1.75 provided a coherent picture.

### Between-phase comparison

Interestingly, we found a significant interaction between the phase (learning or transfer) and transfer feedback (without/with) on correct choice rate (*F*(1, 792) = 82.30, *p* < .0001, 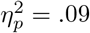). This interaction is shown in **Figure 3** and reflects the fact that while adding transfer feedback information had a significant effect on transfer performance (*F*(1, 792) = 137.18, *p* < .0001, 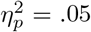, **Figure 3A-B**), it was not sufficient to outperform learning performance (with transfer feedback: learning performance 0.69 ± 0.16 vs transfer performance 0.68 ± 0.15, *t*(399) = 0.89, *p* = .38, *d* = 0.04, **Figure 3B**). Finally, close inspection of the learning curves revealed that in experiments where feedback was not provided in the transfer phase (E1, E3, E5 and E7), choice rates (and therefore option preferences) were stationary (**Figure 3A** and **Figure 3B**). This observation rules out the possibility that reduced performance in the transfer phase was induced by progressively forgetting the values of the options (in which case we should have observed a non-stationary and decreasing correct response rate).

**Figure 3:**
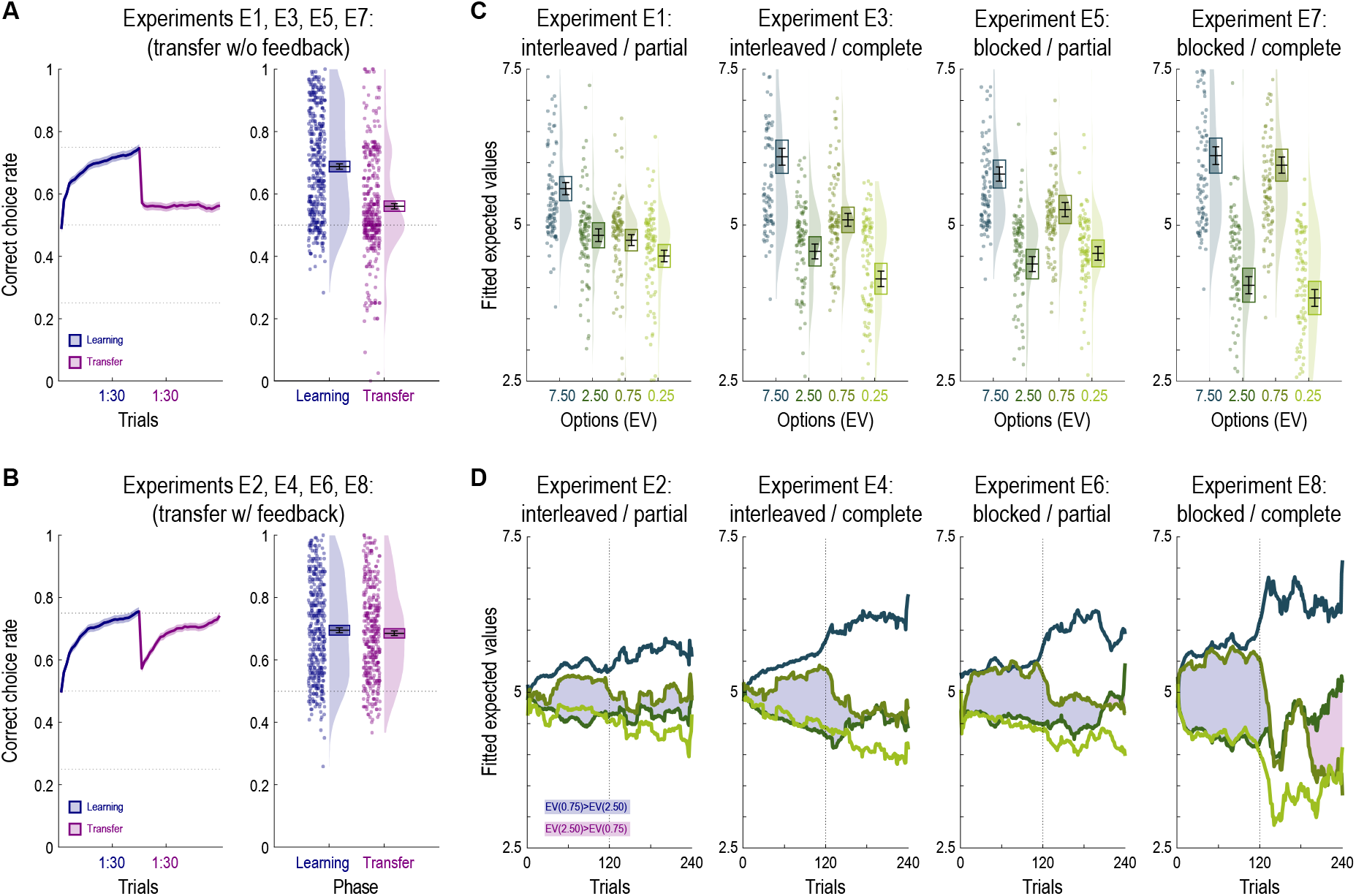
Learning versus transfer comparison and inferred option values. **(A-B)** Average response rate in the learning (blue) and transfer (pink) phase for experiments without **(A)** and with **(B)** trial-by-trial transfer feedback. Leftmost panel: learning curves; rightmost panel: average across all trials. **(C)** Average inferred option values for the experiments without trial-by-trial transfer feedback. **(D)** Trial-by-trial inferred option values for the experiments with trial-by-trial transfer feedback. In all panels: points indicate individual average, areas indicate probability density function, boxes indicate 95% confidence interval and errors bars indicate s.e.m.

In conclusion, comparison between the learning and the transfer phase reveals two inter-related and intriguing facts: i) despite the fact that the transfer phase happens immediately after an extensive learning phase, performance is, if anything, lower compared to the learning phase; ii) factors that improve performance (by intrinsically or extrinsically reducing outcome uncertainty) in the learning phase have either no (feedback information) or a negative (task structure) impact on the transfer phase performance.

### Inferred option values

To visualize and quantify how much observed choices deviate from the experimentally determined true option values, we optimized the four possible option values as free parameters. More precisely, we initialized each subjective value at their true value (we labeled the four possible expected values as follows: EV_7.5_, EV_2.5_, EV_0.75_, and EV_0.25_), and optimized these values by gradient descent in order to maximize the likelihood of observing participants’ choices using the logistic function (for, say, options EV_2.5_ and EV_0.75_):

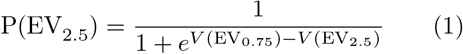

So that, if a participant chose indifferently between the EV_2.5_ and the EV_0.75_ options, their fitted values would be very similar: V(EV_2.5_) ≈ V(EV_0.75_). Conversely, a participant with a sharp (optimal) preference for EV_2.5_ over EV_0.75_ would lead different fitted values: V(EV_2.5_) > V(EV_0.75_). In a first step, in the experiments where feedback was not provided in the transfer phase (E1, E3, E5 and E7), we optimized a set of subjective values per participant.

Consistent with the correct choice rate results described above, we found a value inversion of the 2 intermediary options (EV_2.5_4.46 ± 1.2, EV_0.75_5.26 ± 1.2, *t*(399) = −7.82, *p* < .0001, *d* = −0.67), which were paired in the ΔEV=1.75 context (**Figure 3C**). The differential was also strongly modulated across experiments (*F*(3, 396) = 18.9, *p* < .0001, 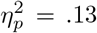, **Figure 3C**) and reached its highest value in E7 (complete feedback and blocked design).

As a second step, in the experiments where feedback was provided in the transfer phase (E2, E4, E6 and E8), we optimized a set of subjective values per trial. This fit allows us to estimate the trial-by-trial evolution of the subjective values over task time. The results of this analysis clearly show that suboptimal preferences progressively arise during the learning phase and disappear during the transfer phase (**Figure 3D**). However, the suboptimal preference was completely corrected only in E8 (complete feedback, blocked design) by the end of the transfer phase.

The analysis of inferred option values clearly confirms that participants’ choices do not follow the true underlying objective monotonic ordering of the option values. Furthermore, it also clearly illustrates that in choice contexts that are supposed to reduce the uncertainty about the option values (complete feedback, blocked design), the deviation from monotonic ordering is paradoxically even greater.

### Computational formalization of the behavioral results

To formalize context-dependent reinforcement learning and account for the behavioral results, we designed a modified version of a standard model, where option-dependent Q-values are learnt from a relative reward term. At each trial *t*, the relative reward *R_REL,t_* is calculated as follows:

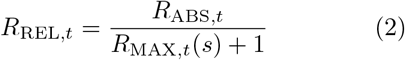

where *s* is the decision context (i.e., a combination of options) and *R*_MAX_ is a context-dependent variable, initialized to 0 and updated at each trial t if the outcome is greater than its current value. As such, *R*_MAX_ will converge to the maximum outcome value in each decision context, which in our task is either 1pt or 10pt. In the first trial *R*_REL_ = *R*_ABS_ (because *R*_MAX, *t*_(*s*) = 0), and in later trials it is progressively normalized between 0 and 1 as the range value *R*_MAX_(*s*) converges to its true value. We refer to this model as the RELATIVE^1^ model and we compared it to a benchmark model (ABSOLUTE) which updates option values based the absolute reward values (note that the ABSOLUTE is nested within the RELATIVE model). For each model, we estimated the optimal free parameters by likelihood maximization. We used the out-of-sample likelihood to compare goodness-of-fit and parsimony of the different models. To calculate the out-of-sample likelihood in the learning phase, the optimization was performed on half of the trials (one ΔEV=5.0 and one ΔEV=0.5 decision context) in the learning phase and the best fitting parameters in this first set were used to predict choices in the remaining half of trials. In the learning phase, we found that the RELATIVE model significantly outperformed the ABSOLUTE model (out-of-sample LL_REL_ vs LL_ABS_, *t*(799) = 6.89, *p* < .0001, d = 0.24). To calculate the out-of-sample likelihood in the transfer phase, the optimization was performed on all trials of the learning phase and the best fitting parameters in the learning phase were used to predict choices in the transfer phase. Thus, the resulting likelihood is not only out-of-sample, but also cross-learning phase. This analysis revealed that the RELATIVE model outperformed the ABSOLUTE model (out-of-sample LL_REL_ vs LL_ABS_, *t*(799) = 8.56, *p* < .0001, *d* = 0.30).

To study the behaviors of our computational model and confirm the behavioral reasons underlying the out-of-sample likelihood results, we simulated the two models (using the individual best fitting parameters) [18]. In the learning phase, only the RELATIVE model managed to reproduce the observed correct choice rate. Specifically, the ABSOLUTE model predicts very poor performance in the ΔEV=0.5 context (ABS vs. data, *t*(799) = −16.90, *p* < .0001, *d* = 0.60, REL vs. data, *t*(799) = −1.79, *p* = .07, *d* = −0.06, **Figure 4A**). In the transfer phase, and particularly in the ΔEV=1.75 context, only the RELATIVE model manages to account for the observed correct choice rate, while the ABSOLUTE model fails (ABS vs. data, *t*(799) = 13.20, *p* < .0001, *d* = 0.47, REL vs. data, *t*(799) = 0.36, *p* = .72, *d* = 0.01, **Figure 4C-D**). In general, the ABSOLUTE model tends to overestimate the correct choice rate in the transfer phase.

**Figure 4:**
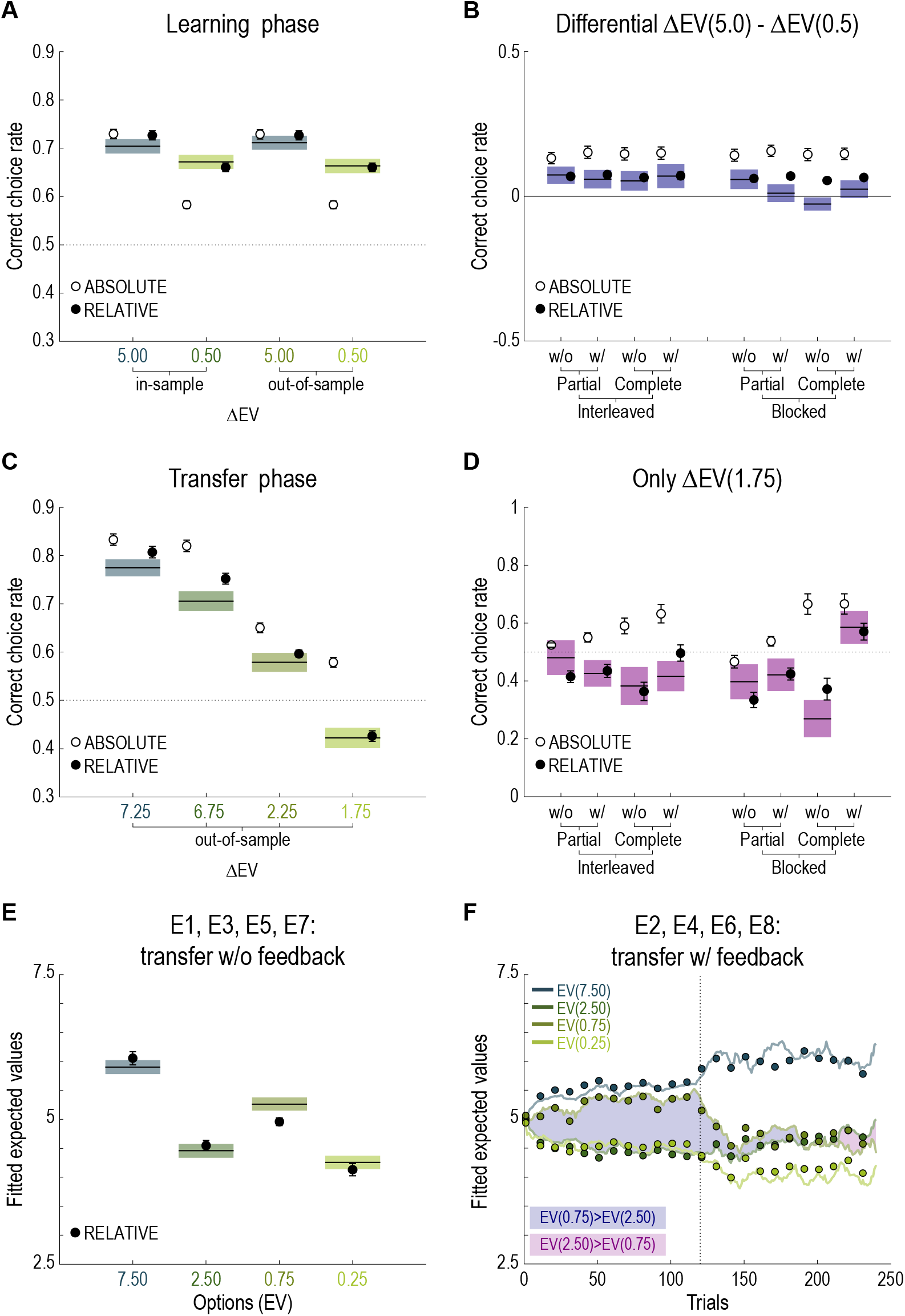
Model comparison. Model simulations of ABSOLUTE (white) and RELATIVE (black) models over the behavioral data (mean and 95% confidence interval) in each context. **(A)** Simulated data in the learning phase were obtained with the parameters fitted in half the contexts (ΔEV=5.0 and ΔEV=0.5) of the learning phase (in-sample and out-of-sample predictions). **(B)** Data and simulations of the differential between high magnitude (ΔEV=5.0) and low magnitude (ΔEV=0.5) contexts. **(C)** Simulated data in the transfer phase were obtained with the parameters fitted in all the contexts of the learning phase (out-of-sample predictions). **(D)** Data and simulations in the context ΔEV=1.75 only. **(E)** Average inferred option values for the behavioral data and simulated data (black dots: RELATIVE model only) for the experiments without trial-by-trial transfer feedback. **(F)** Trial-by-trial inferred option values for the behavioral data and simulated data (colored dots: RELATIVE model only) for the experiments with trial-by-trial transfer feedback, where curves indicate trial-by-trial fit of each inferred option value^9^, and colored dots indicate RELATIVE model simulations.

In addition to these behavioral predictions, we performed a learning-free optimization on RELATIVE model simulations. Not only is the RELATIVE model able to capture the value inversion that we observed in the data, as well as the estimated options (REL vs data, *t*(799) = 1.55, *p* = .12, *d* = 0.06, **Figure 4E**), but it is also able to predict its dynamic emergence and its trial-by-trial evolution (**Figure 4F**).

### Ruling out habit formation

One of the distinguishing behavioral signatures of the RELATIVE model compared the ABSOLUTE one is the preference for the suboptimal option in the ΔEV=1.75 context. Since the optimal option in the ΔEV=1.75 context is not often chosen during the learning phase (where it is locally suboptimal), it could be argued that this result arises from taking decisions based on a weighted average between their absolute values and past choice propensity (a sort of habituation or choice trace). To rule out this interpretation, we fitted and simulated a version of a HABIT model, which takes decisions based on a weighted sum of the absolute Q-values and a habitual choice trace [17,19]. The habitual choice trace component is updated with an additional learning rate parameter that gives a bonus to the selected action. Decisions are taken comparing option-specific decision-weights *D_t_*:

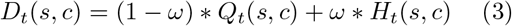

where at each trial *t*, state *s* and chosen option *c, ω* is the arbiter, *Q* is the absolute Q-value *H* is the habitual choice trace component. The weight *ω* is fitted as an additional parameter (for ω = 0 the model reduces to the ABSOLUTE model) and governs the relative influence of each controller.

We found that the HABIT model, similarly to the ABSOLUTE model, fails to perfectly match the participants’ behavior, especially in the ΔEV=0.5 and ΔEV=1.75 contexts (**Figure 5A**). Indeed, in the learning phase, the addition of a habitual component is not enough to cope for the difference in option values, and therefore the model simulations in the transfer phase fail to match the observed choice pattern (**Figure 5B**). This is because the HABIT model encodes values on an absolute scale and does not manage to develop a strong preference for the correct response in the ΔEV=0.5 context, in the first place (**Figure 5A**). Thus, it does not carry a choice trace strong enough to overcome the absolute value of the correct response in the ΔEV=1.75 context (**Figure 5B**, **Supp. Figure 2 A-B**). To summarize, a model assuming absolute value encoding coupled with a habitual component could not fully explain observed choices in both the learning and transfer phase.

**Figure 5:**
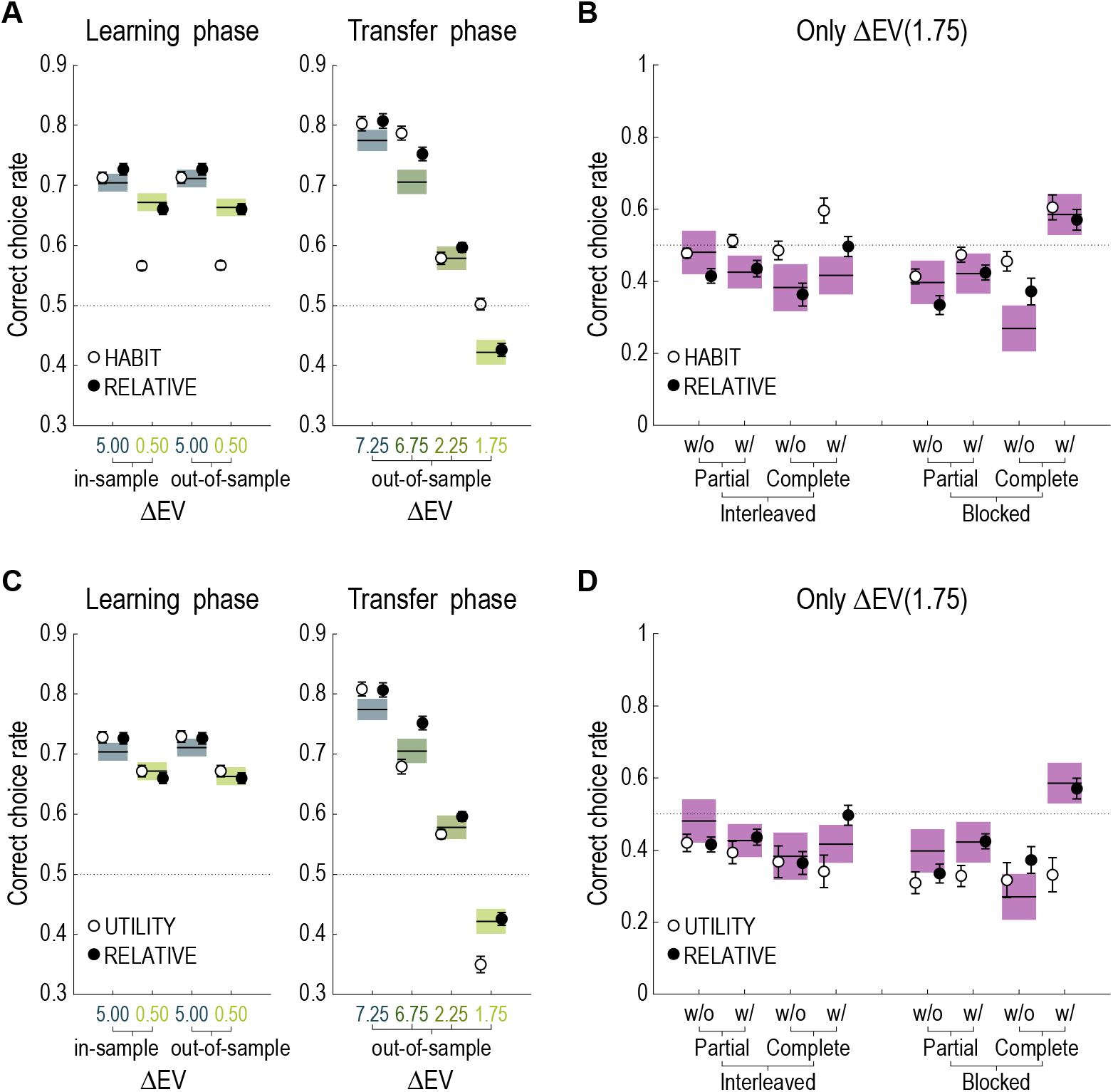
Ruling out the HABIT and UTILITY models. Model simulations of HABIT, resp. UTILITY (white) and RELATIVE (black) models over the behavioral data (mean and 95% confidence interval) in each context. **(A, C)** Simulated data in the learning phase were obtained with the parameters fitted in half the contexts (ΔEV=5.0 and ΔEV=0.5) of the learning phase (in-sample and out-of-sample predictions). Simulated data in the transfer phase were obtained with the parameters fitted in all the contexts of the learning phase (out-of-sample predictions). **(B, D)** Data and simulations in the context ΔEV=1.75 only.

### Ruling out marginally decreasing utility

One of the distinguishing behavioral signatures of the RELATIVE model is that it predicts very similar correct choice rates in the ΔEV=5.00 and the ΔEV=0.50 contexts compared to the behavioral data, while both the ABSOLUTE and the HABIT predict a huge drop in performance in the ΔEV=0.50 that directly stems from its small difference in expected value. It could be argued that this result arises from the fact that expected utilities (and not expected values) are learned in our task. Specifically, a marginally decreasing utility parameter would blunt differences in outcome magnitudes and would suppose that choices are made comparing outcome probabilities. The process could also explain the preference for the suboptimal option in the ΔEV=1.75 context, since the optimal option in the ΔEV=1.75 context is rewarded (10pt) only 25% of the time, while the suboptimal option is rewarded (1pt) 75% of the time. To rule out this interpretation, we fitted and simulated a UTILITY model, which updates Q-value based reward utilities calculated from absolute reward as follows:

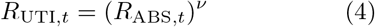

where the exponent *ν* is the utility parameter (0 < *ν* < 1, for *ν* =1 the model reduces to the ABSOLUTE model).

We found that the UTILITY model, similarly to the RELATIVE model, captures quite well the participants’ behavior, in the ΔEV=0.5 context (**Figure 5C**). However, concerning the transfer phase (especially the ΔEV=1.75 context), it fails to capture the observed pattern (**Figure 5C-D**). Additional analyses suggest that this is specifically driven by the experiments where the feedback was provided during the transfer phase (**Figure 5D**). Indeed, the static nature of the UTILITY fails to match the fact that the preferences in the ΔEV=1.75 context can be reversed by providing complete feedback (**Supp. Figure 2 CD**). To summarize, a model assuming diminishing marginal utilities could not fully explain observed choices in the transfer phase.

### Sub-optimality of range-adaptation in our task

The RELATIVE model is computationally more complex compared to the ABSOLUTE model, as it presents an additional internal variable (*R*_MAX_), which is learnt with a dedicated parameter. Here we wanted to assess whether this additional computational complexity really paid off in our task.

We split the participants according to the sign of out-of-sample likelihood difference between the RELATIVE and the ABSOLUTE model: if positive, the RELATIVE model better explains the participants’ data (REL>ABS), if negative, the ABSOLUTE model does (ABS>REL). Reflecting our overall model comparison result, we found much more subject in the REL>ABS, compared to the ABS>REL category (N=545 vs N=255).

We found no main effect of winning model on overall (both phases) performance (*F*(1,798) = 0.03, *p* = .87, 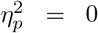). Interestingly, we found that while RELATIVE encoding is beneficial and allows for better performance in the learning phase, it leads to worst performance in the transfer phase (*F*(1, 798) = 187.3, *p* < .0001, 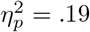, **Figure 6A**). As in the principle of communicating vessels, increasing/decreasing performance in the learning phase seems to correspond to lowered/enhanced performance in the transfer phase. A second question is whether overall in our study, behaving as a RELATIVE model revealed economically advantageous. To answer this question, we compared the final monetary payoff in the real data and following the simulations using the subject level best fitting parameters. Consistently with the task design, we found that the monetary outcome was higher in the transfer phase than in the learning phase (transfer gains M = 2.16 ± 0.54, learning gains M = 1.99 ± 0.35, *t*(799) = 8.71, *p* < .0001, *d* = 0.31). Crucially, we found that the simulation of the RELATIVE model induces significantly lower monetary earnings (ABS vs REL, *t*(799) = 19.39, *p* < .0001, *d* = 0.69, **Figure 6B**). This result indicates that despite being locally adaptive (in the learning phase), in our task range adaptation is economically disadvantageous, thus supporting the idea that it is the consequence of automatic, uncontrolled, process.

**Figure 6:**
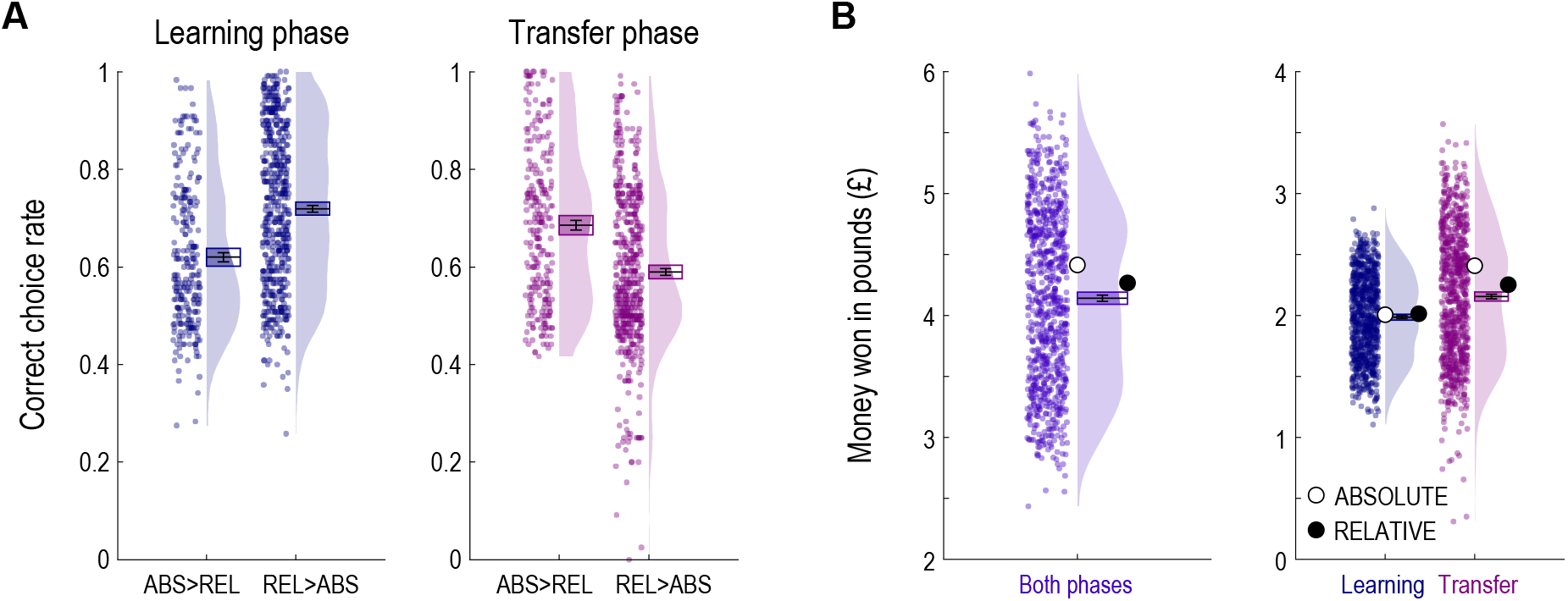
Ruling out the HABIT and UTILITY models. Model simulations of HABIT, resp. UTILITY (white) and RELATIVE (black) models over the behavioral data (mean and 95% confidence interval) in each context. **(A, C)** Simulated data in the learning phase were obtained with the parameters fitted in half the contexts (ΔEV=5.0 and ΔEV=0.5) of the learning phase (in-sample and out-of-sample predictions). Simulated data in the transfer phase were obtained with the parameters fitted in all the contexts of the learning phase (out-of-sample predictions). **(B, D)** Data and simulations in the context ΔEV=1.75 only.

## Discussion

In the present paper we investigate contextdependent reinforcement learning, more specifically range adaptation, in a large cohort of human participants tested online over eight different variants of a behavioral task. Building on previous studies of context-dependent learning, the core idea of the task is to juxtapose an initial learning phase with fixed pairs of options (featuring either small or big outcomes) to a subsequent transfer phase where options are rearranged in new pairs (mixing up small and big outcomes [6, 7, 10]). In some experiments, we intrinsically reduced option value uncertainty by providing complete feedback. In some experiments, we extrinsically modulated option value uncertainty by clustering in time the trials of a given contexts. Finally, in some experiments, feedback was also provided in the transfer phase.

### Behavioral findings

Expectedly, correct choice rate in the learning phase was higher when the feedback was complete and the design blocked, which indicates that participants do pay attention to the outcome of the forgone option and benefit from facilitating option identification and value retrievals findings, we found that, overall, correct response rate was slightly but significantly higher in the big magnitude contexts (ΔEV=5.0), but the difference was much smaller compared to what one would expect assuming absolute value learning and representation (as showed by the ABSOLUTE model simulations [6]): a pattern consistent with a partial range adaptation. The outcome magnitude-induced difference in correct choice rate was significantly smaller and not different from zero in blocked experiments (full adaptation), thus providing a first hint that reducing task difficulty increases range adaptation. Despite learning phase performance being fully consistent with our hypothesis, the crucial evidence comes from the results of the transfer phase. First, overall correct response rate pattern in the transfer phase did not follow that of the learning phase. Complete feedback and blocked design factors have no direct beneficial effect on transfer phase performance. In fact, if anything, the worst possible transfer phase performance was obtained in a complete feedback and blocked experiment. This was particularly striking in the ΔEV=1.75 condition, where participants significantly preferred the suboptimal option and, again, worst score was obtained in a complete feedback and blocked design experiment. Second, we ruled out that the comparably low performance in the transfer phase was due to having forgotten the value of the options. Indeed, since the transfer phase is, by definition, after the learning phase, although very unlikely (the two phases were only few seconds apart), it is conceivable that a drop in performance is due to the progressive forgetting of the option values. Two features of the correct choice rate curves allowed to reject this interpretation: i) correct choice rate abruptly decreases just after the learning test; ii) when feedback is not provided the choice rate remains perfectly stable with no sign of regression to chance level. On the other side, i.e., when feedback was provided in the transfer phase, the correct choice rate increased to reach (on average) the level research at the end of the learning phase. The results are therefore consistent with the idea that in the transfer phase, participants express context-dependent option values acquired during the learning phase, which entails a first counter-intuitive phenomenon: even if the transfer phase is performed immediately after the learning phase, the correct choice rate drops. This is due to the rearrangement of the options in new choice contexts, where options that were previously optimal solutions (in the small magnitude contexts) become suboptimal solutions. We also observed a second counter-intuitive phenomenon: factors that increase performance during the learning phase by extrinsically and intrinsically reducing value uncertainly, paradoxically further undermined transfer phase correct choice rate. The conclusions based on these behavioral observations were confirmed by inferring the most plausible option values based on the observed choices, where we could compare the objective ranking of the options to their subjective estimation. The only experiment where we observed an almost monotonic ranking was the partial feedback / interleaved experiment, even if we observed no significant different between the EV=2.5 and the EV=0.75 options. In all the other experiments, the EV=0.75 option was valued more compared to the EV=2.5 option, with the highest difference observed in the complete feedback / blocked design. Thus, in striking opposition with the almost universally shared intuition that reducing uncertainty should lead to more accurate subjective estimation, here we present a rare instance where the opposite is true.

### Computational modeling

The observed behavioral results were perfectly captured by a parsimonious model (the RELATIVE model) that instantiated a dynamic range adaptation process. Specifically, the RELATIVE model learns in parallel a context-dependent variable (*R*_MAX_) that is used to normalize the outcomes. The *R*_MAX_ is learnt incrementally and its speed determines the extent of the normalization, leading to partial or full range adaptation as a function of the contextual learning rate. Developing a new model was necessary, as previous models of context-dependent reinforcement learning did not include range adaptation and focused on different dimensions of context-dependence (reference point-centering and outcome comparison [7, 8]). The model represents also an improvement over a previous study where we instantiated partial arrange adaptation assuming a perfect and innate knowledge about the context-level variables and a static hybridization process [6].

One limitation of the RELATIVE model is that it approximates the range to the value of its maximum outcome. Of course, this is well suited for our task, where the minimum possible outcome is always zero. This is of course an unrealistic assumption as, both in the lab and in real life, outcome ranges change in terms of both maximum and minimum outcomes. However, this limitation can be easily overcome assuming that our model is a special case of a model that tracks both maximum and the minimum outcome:

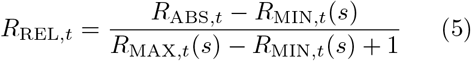

As *R*_min_ is initialized at zero and never changes, in our task this model can be reduced the one presented in equation (2). Another limitation is that in present formulation *R*_MAX_ can only grow. Again, this is a feature that is well suited for our task, but may not correspond to many other laboratory-based and real-life situations, where *R*_MAX_ can drift over time. This limitation could be overcome by assuming *R*_MAX_ is also updated at a smaller rate when the observed outcome is smaller than the current *R*_MAX_. Finally, we note that our model could also be seen as special case of a divisive normalization process (temporal normalization [20]). Future research contrasting different outcome ranges and three-options learning task is required to determine whether the functional form of context-dependent normalization follows a range adapting or divisive scheme [21].

We compared and ruled out another plausible computational interpretation derived from prominent psychological theory [22,23]. First, we considered a habit formation model [17]. We reasoned that our transfer phase results (and particularly the value inversion in the ΔEV=1.75 context) could derive from the participants choosing based on a weighted average between absolute values and past choice propensities. In fact, the suboptimal option in the ΔEV=1.75 context (EV=0.75) was chosen more frequently than the optimal option (EV=2.5) in the learning phase. However, model simulations showed that the HABIT model was not capable to explain the observed pattern. In fact, in the learning phase, the HABIT model, just like the ABSOLUTE model, did not develop a preference for the EV=0.75 option strong enough to generate a habitual trace sufficient to explain the transfer phase pattern. Beyond model simulation comparisons, we believe that this interpretation could have been rejected based on a priori arguments. The HABIT model can be conceived as a way to model habitual behavior, i.e., responses automatically triggered by stimulus-action associations. However, both in real life and laboratory experiments, habits have been shown to be acquired over time scales (days, months, year) order of magnitudes bigger compared to the timeframe of our tasks [24,25]. Indeed, it is even debatable whether in our task participants developed even a sense of familiarity toward the (never seen before) abstract cues we used as stimuli. The HABIT model can also be conceived as a way to model choice hysteresis, sometimes referred to as choice repetition of perseveration bias, that could arise from a form of sensory-motor facilitation, where recently performance actions become facilitated [19,26]. However, the screen position of the stimuli was randomized in a trial-by-trial basis; most of the experiments involved inter-leaved design, thus precluding any strong role for sensory-motor facilitation-induced choice inertia.

We compared and ruled out a plausible computational interpretation derived from economic theory [27]. Since Bernoulli (1700-1782), risk aversion is explained by assuming diminishing marginal utility of objective outcomes [28]. At the limit, if diminishing marginal utility was applied in our case, the utility of 10 points could be perceived as the utility of 1 point. In this extreme scenario, choices would be only based on the comparison between the outcome probabilities. This could explain most aspects of the choice pattern. Indeed, the UTILITY model did a much better job compared to HABIT model. However, compared to the RANGE model, it failed to reproduce the observed behavior of the experiments were feedback was provided in the transfer phase. This naturally results from the fact that the model assumes diminishing marginal utility as being a static property of the model and therefore cannot account for experience-dependent correction of context-dependent biases. However, also in this case, a priori considerations could have ruled out the UTILITY interpretation. Our experiment involves stakes small enough to make diminishing marginal utility not reasonable. M. Rabin provides a full treatment of this issue, and shows that the explaining diminishing marginal utility for small stakes (as those used in the laboratory) leads to extremely unlikely prediction, such as turning down gambles with infinite positive expected values [16]. Indeed, following H. Markowitz intuition, most realistic models of the utility function suppose risk neutrality (or risk seeking) for small gains [29].

### Conclusions

To conclude, we demonstrated that in humans, reinforcement learning values are learnt in a contextdependent manner that is compatible with a range adaptation process [30]. Specifically, we tested the possibility that this results from the way outcome information is automatically processed to achieve adaptive coding [31], by showing that the lower outcome uncertainty, the fuller range adaptation. This leads to a paradoxical result: reducing task difficulty can, in some occasions, decrease choice optimality. This surprising result can be understood by coming back to a perceptual analogy. Going into a dark room forces us to adapt our retinal response to dark, so that when we go back into a light condition we do not see very well. The longer we are exposed to dim light, the stronger the effect when we go back to normal. Our findings fit in the debate aimed at deciding whether the computational processes leading to suboptimal decisions have to be considered flaws or feature of human cognition [32,33]. Range adapting reinforcement learning is clearly adaptive in the learning phase. We could hypothesize that the situations in which the process is adaptive are more frequent in real life. In other terms the performance of the system has to be evaluated as a function of the tasks it has been selected to solve. It is true that we may be hit by a bus when we exit a dark room because we do not see well, but on average, the benefit of a sharper perception in a dark room is big enough to compensate for the (rare) event of a bus waiting for us outside the dark room. Ultimately, whether context-dependent reinforcement learning should be considered a flaw or a desirable feature of human cognition would be determined by how frequent are the situations where it is adaptive (as in the learning phase) compared to those where it is maladaptive (as in the transfer phase). While our study does not settle this issue, by showing that range adaptation in a task where it is economically disadvantageous, our findings suggest that this process may, at least in some circumstances, contribute to suboptimal choices.

## Materials and methods

### Participants

For the laboratory experiment, we recruited 40 participants (28 females, aged 24.28±3.05 years) via Internet advertising in a local mailing-list dedicated to cognitive science-related activities.

For the online experiments, we recruited 8×100 participants (414 females, aged 30.06±10.10 years) from the Prolific platform (www.prolific.co). The research was carried out following the principles and guidelines for experiments including human participants provided in the declaration of Helsinki (1964, revised in 2013). The Inserm Ethical Review Committee / IRB00003888 approved the study on November 13th, 2018 and subjects were provided written informed consent prior to their inclusion. To sustain motivation throughout the experiment, subjects were given a bonus depending on the number of points won in the experiment (average money won in pounds: 4.14±0.72, average performance against chance: M = 0.65±0.13, *t*(799) = 33.91, *p* < 0.0001). A laboratory-based experiment was originally performed (N=40) to ascertain that online testing would not significantly affecting the main conclusions. The results are presented in the **Supplementary Materials**.

### Behavioral tasks

Participants performed an online version of a probabilistic instrumental learning task adapted from previous studies [6]. After checking the consent form, participants received written instructions explaining how the task worked and that their final payoff would be affected by their choices in the task. During the instructions the possible outcomes in points (0pt, 1pt and 10pt) were explicitly showed as well as their conversion rate (1pt=£0.005). The instructions were followed by a short training session of 12 trials aiming at familiarize the participants with the response modalities. Participants could repeat the training session up to two times and then started the actual experiment.

In our task, options were materialized by abstract stimuli (cues) taken from randomly generated identicons, colored such that the subjective hue and saturation were very similar according to the HSLUV color scheme (www.hsluv.org).On each trial, two cues were presented on both sides of the screen. The side in which a given cue was presented was pseudo-randomized, such that a given cue was presented an equal number of times in the left and the right. Subjects were required to select between the two cues by clicking on the cue. The choice window was self-paced. A brief delay after the choice was recorded (500 ms), the outcome was displayed for 1000 ms. There was no fixation screen between trials. The average reaction time was 1.36±0.04 seconds (median: 1.16), the average experiment completion time was 325.24±8.39 seconds (median: 277.30).

As in previous studies, the full task consisted in one learning phase followed by a transfer phase [6–8, 34]. During the learning phase, cues appeared in 4 fixed pairs. Each pair was presented 30 times, leading to a total of 120 trials. Within each pair, the two cues were associated to a zero and a non-zero outcome with reciprocal probabilities (0.75/0.25 and 0.25/0.75). At the end of the trial, the cues disappeared and the selected one was replaced by the outcome (“10”, “1”, or “0”) (**Figure 1A**). In Experiments 3, 4, 7 and 8, the outcome corresponding to the forgone option (sometimes referred to as the counterfactual outcome) was also displayed (**Figure 1C**). Once they had completed the learning phase, participants were displayed with the total points earned and their monetary equivalent.

During the transfer phase after the learning phase, the pairs of cues were rearranged into 4 new pairs. The probability of obtaining a specific outcome remained the same for each cue (**Figure 1B**). Each new pair was presented 30 times, leading to a total of 120 trials. Before the beginning of the transfer phase, participants were explained that they would be presented with the same cues, only that the pairs would not have been necessarily displayed together before. In order to prevent explicit memorizing strategies, participants were not informed that they would have to perform a transfer phase until the end of the learning phase. After making a choice, the cues disappeared. In experiments 1, 3, 5 and 7, participants were not informed of the outcome of the choice on a trial-by-trial basis and the next trial began after 500ms. This was specified in the instruction phase. In experiments 2, 4, 6 and 8, participants were informed about the result of their choices in a trial-by-trial basis and the outcome was presented for 1000ms. In all experiments they were informed about the total points earned at the end of the transfer phase. In addition to the presence / absence of feedback, experiments differed in two other factors. Feedback information could be either partial (experiments 1, 2, 5, 6) or complete, (experiments 3, 4, 7, 8; meaning the outcome of the forgone option was also showed). When the transfer phase included feedback, the information factor was the same as in the learning phase. Trial structure was also manipulated, such as in some experiments (5, 6, 7, 8) all trials of a given choice context were clustered (‘‘blocked”), and in the remaining experiments (1, 2, 3, 4) they were interleaved (**Figure 1C**).

### Analyses

#### Behavioral analyses

The main dependent variable was the correct choice rate, i.e., choices directed toward the option with the highest expected value. Statistical effects were assessed using multiple-way repeated measures ANOVAs with choice context (labeled in the manuscript by their difference in expected values: ΔEV) as within-subject factor, and feedback information, feedback in the transfer phase and task structure as between-subjects factors. Post-hoc tests were performed using one-sample and two-sample t-tests for respectively within- and between-experiment comparisons. To assess overall performance, additional one sample t-tests were performed against chance level (0.5). We report the t-statistic, p-value, and Cohen’s d to estimate effect size (two-sample t-test only). Given the large sample size (N=800), central limit theorem allows us to assume normal distribution of our overall performance data and to apply properties of normal distribution in our statistical analyses, as well as sphericity hypotheses. Concerning ANOVA analyses, we report the uncorrected statistical, as well as Huynh-Feldt correction for repeated measures ANOVA when applicable [35], F-statistic, p-value, partial eta-squared 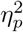 and generalized eta-squared *η*^2^ (when Huynh-Feldt correction is applied) to estimate effect size. All statistical analyses were performed using Matlab (www.mathworks.com) and R (www.r-project.org). For visual purposes, learning curves were smoothed using a moving average filter (span of 5 in Matlab’s smooth function).

### Models

We analyzed our data with variation of simple associative learning models [36, 37]. The goal of all models is to estimate in each choice context (or state) the expected reward (R) of each option and pick the one that maximizes this expected reward R.

At trial t, option values of the current context s are updated with the delta rule:

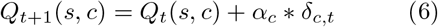

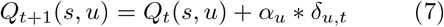

where *α_c_* is the learning rate for the chosen (*c*) option and *α_u_* the learning rate for the unchosen (*u*) option, i.e., the counterfactual learning rate. *δ_c_* and *δ_u_* are prediction error terms calculated as follows:

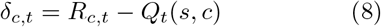

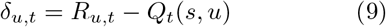

*δ_c_* is calculated in both partial and complete feedback contexts and *δ_u_* is calculated in the experiments with complete feedback only.

We modelled participants’ choice behavior using a softmax decision rule representing the probability for a participant to choose one option a over the other option b:

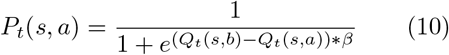

where *β* is the inverse temperature parameter. High temperatures (*β* → 0) cause the action to be all (nearly) equiprobable. Low temperatures (*β* → +∞) cause a greater difference in selection probability for actions that differ in their value estimates [36].

We compared 3 alternative computational models: the ABSOLUTE model, which encodes outcomes in an absolute scale independently of the choice context in which they are presented, the RELATIVE model which tracks the value of the maximum reward in each context and normalizes the actual reward accordingly, rescaling rewards between 0 and 1, and the HABIT model, which integrates action weights into the decision process.

ABSOLUTE model The outcomes are encoded as the participants see them. A positive outcome is encoded as its actual positive value (in points): *R*_ABS, *t*_ ∈ {10,1,0}.

RELATIVE model The outcomes (both chosen and unchosen) are encoded on a context-dependent relative scale. On each trial the relative reward *R*_REL, *t*_ is calculated as follows:

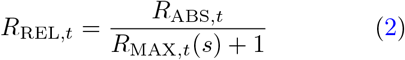

where *s* is the decision context (i.e., a combination of options) and *R*_MAX, *t*_ is a context-dependent variable, initialized to 0 and updated at each trial t if the outcome is greater than its current value:

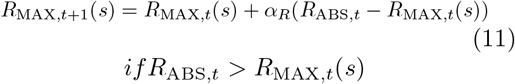

Accordingly, outcomes are progressively normalized so that eventually *R*_REL,*t*_ ∈ [0,1]. The chosen and unchosen option values and prediction errors are updated with the same rules as in the ABSOLUTE model. Note that the RELATIVE model is nested within the ABSOLUTE model (α_R_ = 0).

HABIT model The outcomes are encoded on an absolute scale, but decisions integrate a habitual component [17,19]. To do so, in addition to the Q-values, a habitual (or choice trace) component H is tracked and updated (with a dedicated learning rate parameter) that takes into account the selected action (1 for chosen option, 0 for the unchosen option). The choice is performed with a softmax rule based on decision weights D that integrate Q-values and decision weights H:

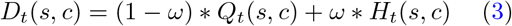

where at each trial *t*, state s and chose option *c*, D is the arbiter, Q is the goal directed component (Q-values matrix), H is the habitual component. The weight *ω* is fitted as an additional parameter and governs the relative weights of values and habits (for *ω* = 0 the model reduces to the ABSOLUTE model).

UTILITY model The outcomes are encoded as an exponentiation of the absolute reward, leading to a curvature of the value function [27]

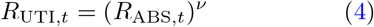

where the exponent *ν* is the utility parameter, with 0 < *ν* < 1 (for *ν* =1 the model reduces to the ABSOLUTE model).

## Supplementary Materials

### Laboratory-based replications and robustness to outliers

To ascertain that online testing would not significantly affecting the main conclusions, a laboratory-based experiment was originally performed. We recruited 40 participants (28 females, aged 24.28±3.05) via Internet advertising in a local mailing-list dedicated to cognitive science-related activities. The experiment design was an exact copy of experiment E2, that is partial feedback information in both the learning phase and the transfer phase, and trials in an interleaved order (**Figure 1**).

In order to characterize learning behavior of participants, we analyzed the correct response rate in both phases, i.e., choices directed toward the most favorable option at each trial. To assess successful learning, we first tested participants’ correct response rate against chance level. We found it to be above chance level in both the learning phase (*t*(39) = 8.88, *p* < .0001, **Supp. Figure 1A**) and the transfer phase (*t*(39) = 5.55, *p* < .0001, **Supp. Figure 1C**), signaling significant instrumental learning effects. We found a significant effect of magnitude in the learning phase (*t*(39) = 2.18, *p* = .036, **Supp. Figure 1B**), and the performance in the ΔEV(1.75) context was significantly below chance level (*t*(39) = −2.43, *p* = .020, **Supp. Figure 1D**), reflecting context-induced irrational preferences. We found that the online experiment replicates the laboratory experiment. Learning performance did not significantly differ between laboratory and online datasets (*t*(138) = 1.67, *p* = .10, *d* = 0.31, **Supp. Figure 1A**), as well as the magnitude effect (*t*(138) = −0.15, *p* = .88, *d* = −0.03, **Supp. Figure 1B**), transfer performance (*t*(138) = 0.62, *p* = .54, *d* = 0.12, **Supp. Figure 1C**) and ΔEV(1.75) performance (*t*(138) = −0.84, *p* = .40, *d* = −0.16, **Supp. Figure 1D**). Of note, we did not find a significant difference between laboratory-based and online-based experiments when looking at the reaction time (*t*(138) = −0.50, *p* = .62, *d* = −0.09), although the control over the measure of reaction times is arguably limited.

This similarity between laboratory- and online-based results supports the definition of online-based experiments as a way to target larger, more diversified populations with reduced administrative and financial costs [38]. The limitations that can be encountered with online-based experiments - such as lower data quality, faster reaction time, lack of engagement from the participants [39, 40] - were not significantly reflected in our data. To further check the robustness of our results, we run analyses excluding subjects presenting unusual task completion time. We approximated participants’ total reaction time over the whole task by a normal distribution and removed outliers at a significance level of 0.05. This led to a removal of 30 participants for the online experiments, and none for the laboratory experiment, leading to a sample of 770 participants for the 8 online experiments and 40 participants for the laboratory experiment. We found that the totality of the results described in the Results section were observable in both datasets, that is, with or without these outliers, thus we decided to include all participants in the statistics reported in the Results section. In conclusion, our results show that the findings are successfully replicated in lab- and online-based experiments. Moreover, our results confirm the findings of previous (more recent) studies comparing both methods, showing that they produce comparable data quality [41, 42].

**Supp. Figure 1.**
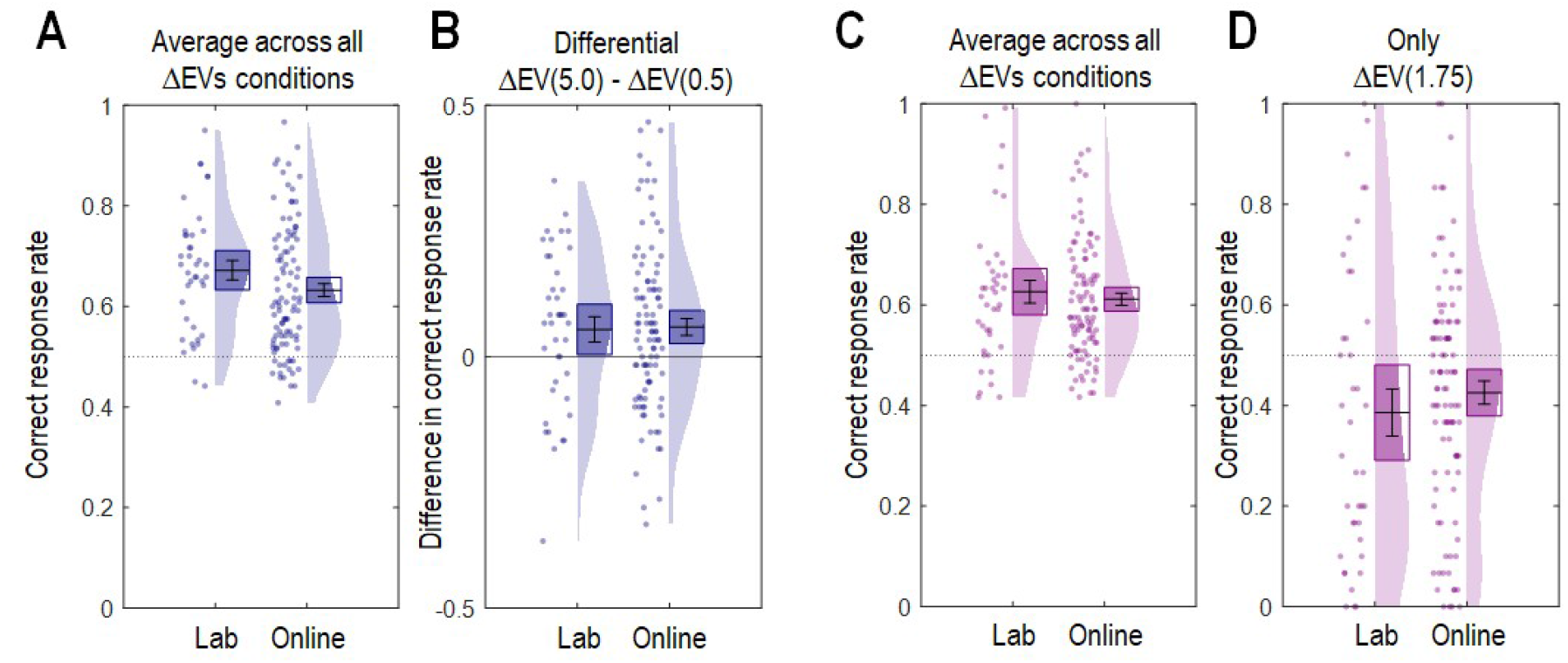
Comparing laboratory and online experiments. **(A)** Average correct response rate in the learning phase per experiment. **(B)** Difference in correct choice rate between the ΔEV=5.0 and the ΔEV=0.5 contexts. **(C)** Average correct response rate in the transfer phase. **(D)** Correct choice rate for the ΔEV=1.75 context only.

**Supp. Figure 2.**
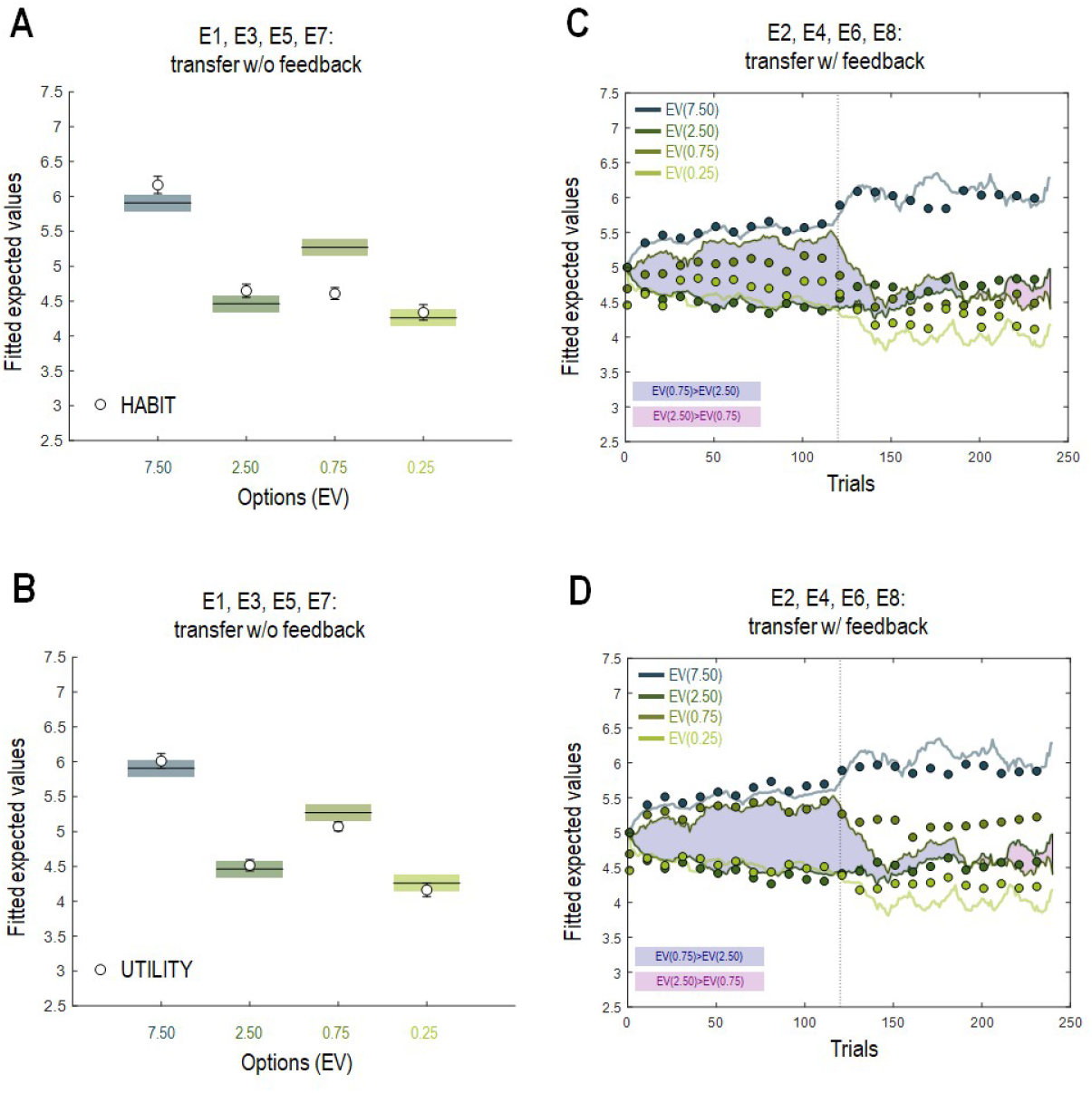
Ruling out habitual learning and marginally decreasing utility. **(A-C)** Average inferred option values for the behavioral data and simulated data for the experiments without trial-by-trial transfer feedback (white dots: HABIT (resp. UTILITY) model). **(B-D)** Trial-by-trial inferred option values for the behavioral data and simulated data for the experiments with trial-by-trial transfer feedback, where curves indicate trial-by-trial fit of each inferred option value, and colored dots indicate HABIT (resp. UTILITY) model simulations.

## Footnotes

1 We note that, while conceptually similar, this model is algorithmically different from previous models also referred to as RELATIVE in previous studies from our and others groups [6–8]

